# Do M-cells contribute significantly in T-wave morphology during normal and arrhythmogenesis conditions like short QT Syndrome?

**DOI:** 10.1101/2020.05.28.121079

**Authors:** Ponnuraj Kirthi Priya, Srinivasan Jayaraman

## Abstract

**Aims:** This paper proposes to explain the mechanism of M-cells, particularly its role in the T-wave generation and its contribution to arrhythmogenesis in short QT syndrome 2 (SQTS2).

**Methods:** A 2D transmural anisotropic ventricular model made up of three principal cell types were developed. Different setups in which: a) entire column of mid-myocardial (mid) cells, b) single island of cells c) two island of cells within the mid-layer d) single island of cells in endocardial (endo)-mid layer were considered as M-cells. These setups are stimulated to explain i) contribution of M-cells in T-wave morphology ii) arrhythmia generation phenomena under SQTS2 heterozygous gene mutation by creating pseudo ECGs from the tissue.

**Results:** Findings infer that setups with an entire layer of M-cells and a higher percentage of epicardial (epi) cells exhibit positive T-waves. Increasing the size of the island in M-cell island setups results in an increased positive T-peak. Placing the M-cell island in the bottom of the mid-layer produced low amplitude T-waves. Further, in two M-cell islands setup, a higher T-wave amplitude was observed when the islands are placed closer than far apart. Moving the M-cell island slightly into the endo layer increases the amplitude of the T-wave. Lastly, on including SQTS2 conditions and pacing with premature beats, an arrhythmia occurs only in those setups containing a layer of M-cells compared to M-cells island setup.

**Conclusion:** These simulation findings paved the way for a better understanding of the M-cells functionality in T-wave morphology as well as promoting arrhythmogenesis under SQTS2 condition.

## 1 Introduction

Ventricular arrhythmias are one of the primary factors responsible for sudden cardiac death. These arrhythmias manifest at the subcellular level due to aberrations in the ionic currents, calcium dynamics or intercellular coupling. Modelling of the ventricle integrating multiscale levels of cardiac function help in dissecting the individual contributions of these different factors. The ventricular myocardium is composed of three principal cells such as endocardial (endo), midmyocardial (mid) and epicardial (epi) cells that vary in their action potential (AP) shape and duration due to the intrinsic differences in their electrophysiological properties. In 1990’s^1^, it was reported that a major contributor to the existence of transmural dispersion of repolarisation (TDR) was the presence of M cells. These cells are found to be localized in the deep subendocardium to midmyocardium in the anterior wall and throughout the wall in the region of RVOT^2^. Later, Näbauer^3^ reported that M-cells are accountable for 30-40 % of the ventricular wall and this relative abundance plays a critical role in a number of electrophysiological phenomena including U-waves and biphasic T-waves.

The hallmark of M cells is its substantial ability to prolong the AP more than that of endo or epi cells at slow heart rates or on exposure to certain drugs like sotalol, quinidine etc. M-cells have longer action potential duration (APD) due to a weaker slow delayed rectifying potassium current (*I_Ks_*) and a stronger late sodium and sodium-calcium exchanger currents. During disease conditions, the mean APD increases and the maximum APD gradient increases markedly. Selective prolongation of APD of M-cells has been known to act as a substrate for promoting conduction block and re-entrant arrhythmia.

For instance, in the intact myocardium, Akar *et al.*, recorded a maximum transmural APD difference of 30-40 ms. In their study, the APD increases from the epicardium to the midmy-cardium reaching a maximum value in the deeper layers and then reducing at the endocardium^4^. Therefore, simulation models adopted the findings of the above study and modelled the M-cells as a uniform bands in the midmyocardial layer in the 1D and 2D. Other experimental studies like that of Glukhov *et. al.*,^5^, found the presence of M-cells in topologies such as isolated islands or clusters with its percentage varying across different regions of the heart.

On the other hand, certain studies dispute the presence of such M-cells with large action potential duration (APD) in the transmural wall and report a smooth transition of APD moving from the endo to epi layers or from the apex to base^6,7^. An amplified APD gradient exists between the epicardial and midmyocardial layers due to a diminished intercellular uncoupling and therefore M cells are more readily detectable under pathologic conditions. Moreover, APD is not directly measured from intact myocardium but are rather estimated from the activation recovery intervals (ARI). Another reason for plunge electrode recordings not being able to pinpoint the location of M-cells accurately is that the anatomic location and topology of M-cells vary considerably in nature.

The existence of M-cells with distinct electrophysiological properties have been confirmed by multiple groups in single cells as well as in tissue wedges and are described henceforth. In the porcine left ventricular wall, M-cells with long APD and characteristic spike and dome configuration were found mostly, but not limited in the midmyocardium^8^. Glukhov *et al.*, optically mapped the APD in left ventricular wedge preparations from failing and non-failing hearts. They observed M-cell islands with longer APD located closer to subendocardium in non-failing human hearts, while in failing human hearts, heterogeneous prolongation of APD in all cells reduced the transmural APD gradients^5^. However, the functional aspects and clinical relevance of M-cells remains controversial. This is in part due to the limitation in measuring techniques to detect M-cells in whole heart^9^. Also, the APD difference measured between M-cells and its neighbouring cells is high in isolated cells compared to intact heart. This can be expected since the intercellular coupling in the tissue would try to homogenize the intrinsic differences in APD between cells. Conrath *et al.*, reported that effective intercellular coupling masks the functionality of M-cells in the heart^10^.

The morphology of the T-wave has been attributed to the differences in the repolarisation times of these different cell types that inherently exist in the ventricular wall as well as the degree of electrotonic coupling^2^. An amplification of this dispersion of repolarisation (DOR) due to genetic factors or effect of drugs has been known to contribute to the development of various cardiac arrhythmia. DOR has been known to exist not only in the transmural layers but also along the apex to base and between the right ventricle and left ventricle. To study how dispersion of APD predominantly contributes to the genesis of T-wave, Keller *et al.*,^11^ created different setups incorporating transmural (TM), apico-basal (AB), interventricular dispersion of APD. They observed that in healthy conditions, the T-waves are usually concordant (same polarity) with the QRS complex in lead II. The simulation results of Keller *et al.*, reported that the best match with clinically observed ECG was obtained with a mix of TM heterogeneity and shorter endo apical APD compared to base APD with M-cells located in the mid-myocardial position. Also, interventricular dispersion led to a notched T-wave while exclusively TM and AB setups led to concordant T-waves.

In case of abnormal pathological conditions, say SQTS, it is characterised by a short QT interval of less than 320 ms^12^ and tall peaked T-waves on the ECG. This shortened action potential distribution of the individual cells in SQTS can increase the risk of atrial and ventricular arrhythmias. Until now, six genetic variants of SQTS have been identified with the first three types of SQTS (1-3) occurring due to gain of function in the genes encoding the different potassium currents while the other three types of SQTS (4-6) are due to a loss of function in the genes encoding the subunits of the L-type calcium current, resulting in a mixed SQT-Brugada syndrome.

SQTS2 occurs due to mutations in the KCNQ1 gene which in combination with KCNE1 encodes the proteins responsible for the slow delayed rectifying potassium current (*I_Ks_*). A g919c substitution in KCNQ1 leads to an aminoacid substitution of valine to leucine on residue 307 (V307L) in the KCNQ1 channel pore helix. This KCNQ1 V307L mutation shifts the voltage dependence of KCNQ1+KCNE1 activation towards more negative voltages and accelerates channel activation, which consequently leads to increased *I_Ks_* current during ventricular repolarisation, thereby reducing the QT interval to 290 ms^13^.

Simulation studies incorporating V307L KCNQ1 SQT2 mutation reproduced the AP shortening of the different cell types. Zhang *et al.*,^14^ incorporated the channel kinetics of the V307L mutation of KCNQ1 subunit of *I_Ks_* using Hodgkin-Huxley (HH) type equations in single cells as well as 1D and 2D transmural tissue. Their study captured important aspects of SQTS such as abbreviated QT interval, increased T-wave amplitude and *T_peak_-T_end_* duration. Further, their simulation result showed that V307L mutation increases transmural heterogeneity of APD, increases the vulnerable window for unidirectional conduction block and therefore facilitates and sustains reentrant arrhythmia. Adeniran *et al.*, simulated the initiation and maintenance of reentrant circuit due to an D172N mutation to Kir 2.1 in SQTS3 condition in 1D, 2D and 3D models^15^. Luo *et al.*, 2017^16^ used these same equations to study the effect of quinidine as a potential drug for terminating SQTS2 under both homozygous and heterozygous mutation conditions of *I_Ks_*. This HH model was unable to capture the slow deactivation of the *I_Ks_* with the V307L KCNQ1 mutation. Hence, Aderinan *et al.*, 2017^13^ incorporated a markov chain (MC) model of *I_Ks_* kinetics for the heterozygous wild type (WT) and homozygous V307L mutant conditions. Their simulations showed that the MC model accurately reproduced AP shortening and reduced effective refractory period in both the mutant conditions which in turn created a reentrant wave sustained at longer duration and higher dominant frequency in 3D human ventricle. Further, partial inhibition of *I_Ks_* successfully terminated the reentry and could be explored as a potential anti-arrhythmic strategy. Multi-scale models have been employed to critically study the ionic mechanisms underlying other variants of SQTS and as well as its proarrhythmic effect in 2D and 3D realistic anatomical geometries. Luo *et al.*,^17^ studied the functional role of M-cells in presence of SQT2 mutation on generating reentrant arrhythmia. Their results show that reentry was easily initiated and sustained for a longer duration in presence of the M-cell island model compared to the M-cell band model. The heterogeneity in APD created by the M-cell plays a vital role in increasing the sustenance of reentry during SQTS2 conditions.

The physical and functional characterization of M-cells have not been completely understood and described until now. This gap provides an opportunity to further explore the effect of position, thickness, and topography of the M-cells as well as its functionality in the formation of T-wave and arrhythmogenesis. These details are difficult to attain through experimental studies, hence this limitation can be bridged by using cardiac in-silico models. Here, the mid-myocardium (mid) layer is considered as an anatomical position in between the endo and epi layer while M-cells are those that have distinct electrophysiological properties. The first part of this paper deals with understanding how the properties of the M-cells: position, size, shape and ionic currents influence the shape of the T-wave. For this different setups are considered in which the M-cells are placed in a two dimensional anisotropic transmural tissue either in the entire mid layer or as islands within the mid layer or at the endo-mid interface. Pseudo ECG are created from each of the different setups and compared with their clinical counterpart to understand which setup produces a realistic match. SQTS2 conditions leads to an abbreviated QT interval, increased *T_peak_* as well as transmural dispersion of repolarisation in the cardiac tissue. Although all the above factors have been postulated to act as a substrate for reentrant arrhythmia, the functional contribution of M-cells in generating such an arrhythmia during SQTS conditions is not very clear. Thus, the second part of this paper deals with understanding how various M-cell topologies and its properties play a role in arrhythmogenesis during SQTS. In those setups that produced an upright T-wave, SQTS2 heterozygous gene mutations are introduced in all cells of the anisotropic transmural tissue to examine which setups would be proarrhythmic.

## 2 Methods

### 2.1 Cell Model

The rise and fall of membrane potential in an individual cell is described by the differential equation below

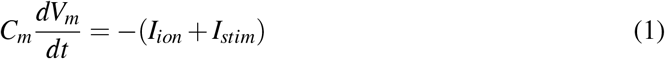

where *V_m_* is the transmembrane potential, t is time, *C_m_* is the membrane capacitance, *I_ion_* is the summation of transmembrane ionic currents developed by Ten Tusscher (TP06)^18^ and *I_stim_* is an external current stimulus. *I_stim_* has an amplitude of 52 *μ*A and is applied for 1 ms in the TP06 model. The differential equations of the ionic parameters are integrated using Rush and Larsen scheme^19^ with a time step of 0.05 ms. The electrophysiological parameters for endo, mid and epi cells were considered from the TP06 model parameters. However, it was found that these original parameters from the TP06 model don’t generate early after depolarisations (EADs), even when *G_Kr_* is reduced to 0. This drawback was overcome by modifying the parameters such as (i) the time constant of the f-gate (*τ_f_*) of the L-type Ca current is decreased by a factor of two and (ii) the conductance of the Ca current (*G_CaL_*) is increased by two fold, in all cell types^20^.

### 2.2 Two dimensional Tissue Model

A transmural section of a 2D ventricular wall having a length of 5 cm and a thickness of 2 cm is considered for evaluation. This section is represented by an array of 250×100 cells in which the space between two cells is assumed to be 0.02 cm. The origin (Cell 1,1) in this array of cells is taken as the bottom leftmost corner. The functionality of gap junctions (GJ) that allow the passage of electric current between cells is modelled using conductive elements. Ohm’s law is used to describe this flow of current between two cells when there exists a difference in membrane voltage between them and it is represented as^21^.

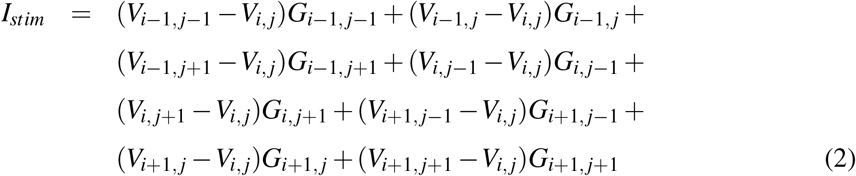

where *I_stim_* is the current used to stimulate cell (i,j) and it is the sum of currents from the eight neighbouring cells. Cells at the corner and perimeter of the array are coupled to 3 and 5 adjacent cells respectively. In the normal myocardial fibres, anisotropy exists with the longitudinal conduction being thrice as fast as the transverse conduction. To incorporate this property in the array, the gap junction conductance (GJC) values along the length and width of the array are set to 4 *μ*S and 0.4 *μ*S respectively which is within the physiological range^22^. These conduction values result in a conduction velocity of 76.9 cm/s and 33.3 cm/s through the length and width of the fibre respectively which is the observed values in experiments of^23^.

### 2.3 Properties of M-cell Layer

In this study transmural tissue was classified into two broad class: Firstly, all the cells in mid layer are considered to be M-cells, and secondly, in which an island of M-cells are considered to be either located completely within the mid layer or partially in the endo layer.

#### 2.3.1 M-cell as an Entire mid Layer

In general, different transmural section set has been reported based on its % of cell in the different layer as its summarised in Table 1. First setup (TM-40)^11^ assumed a fraction of 40% endo, 30% mid and 30% epi with smaller apex APD than the basal APD, and got a good correspondence with the clinically observed ECG, as discussed in the *Introduction* section. Adeniran *et al.*,^24^ had divided the transmural section along the thickness from left to right into 25% endo, 35% mid and 40% epi cells. These percentages were also reported to get an upright T-wave in the ECG and have thus been considered for the second setup (TM-25) along with apex-base variation. Finally TM-30, here transmural section was divided into 30% endo, 30% mid and finally 40% epi cells along with the apex-base variation.

**Table 1:** Percentage of cells in the three setups of M-cell Layer

| Setup | Endo Cells | M/Mid Cells | Epi Cells |
| --- | --- | --- | --- |
| TM-40 | 40 | 30 | 30 |
| TM-25 | 25 | 35 | 40 |
| TM-30 | 30 | 30 | 40 |

Further, properties of different cell layer could be subgrouped based on the ionic parameter variation:

- The conductance of slow delayed rectifying potassium current *G_Ks_* value of endo cells is decreased to 92% of the actual TP06 value so that the APD of endo cells is greater than that of epi cells ^25^
- A graded apex-base variation is introduced in endo cells by increasing the *G_Ks_* of apex to 1.5 times its base value. The values between apex to base are linearly interpolated. In the deep sub-epicardium, a decrease in tissue resistivity has been attributable to a sharp transition in cell orientation, reduced expression of connexin 43 and increased density of collagen in this region

Gima *et al.*,^25^ incorporated this in their simulation model by decreasing the GJC by fivefold at the epi-mid mural junction. Likewise, a fivefold decrease in GJC at the epi-mid mural junction has been included along the whole length for a thickness of 5 cells (Cells 58 to 62) in all the three setups, whereas there is no reduction in GJC at the endo-mid interface. The *G_Ks_* and *G_to_* values of the different cells are given in Table 2.

**Table 2:** Variation in ionic parameters of different cells

| Parameter | Endo Cells | M/Mid Cells | Epi Cells |
| --- | --- | --- | --- |
| $G_{Ks}$ | 0.36064 nS/pF | 0.098 nS/pF | 0.392 nS/pF |
| $G_{to}$ | 0.073 nS/pF | 0.294 nS/pF | 0.294 nS/pF |

#### 2.3.2 M-cell as Island

M-cells have traditionally been defined as those having the largest APD, hence their *G_Ks_* value is taken as 0.098 nS/pF, and other cell’s *G_Ks_* value is 0.392 nS/pF as similar to the TP06 model configuration. These *G_Ks_* values used for different cell types are listed in row 1 of Table 2. However, when these value is retained for epi cells and that of mid cells is reduced to 0.37632 nS/pF, it is observed that a positive T-wave doesn’t occur. Hence, the same *G_Ks_* value is retained for mid cells while that of endo cells is reduced to 0.36064 nS/pF and that of epi cells is increased to 0.588 nS/pF; so that there is a transition in APD among the layers with the epi cells having the smallest APD followed by the mid, and endo cells as reported by^26^. When, the *G_to_* values are considered from the TP06 model parameters, the base has been shown to have a larger APD compared to the apex, however the gradient of variation is small^27^. In this study, we have adapted the Keller *et al.*,^11^ approach to define the *G_Ks_* value as listed in row 2 of Table 3 (i.e *G_Ks_* value in apex that is 1.5 times of that in base).

**Table 3:** *G_Ks_* values of different cell types in different setups

| Setup | Endo Cells (nS/pF) | Mid Cells (nS/pF) | Epi Cells (nS/pF) | M-Cells (nS/pF) |
| --- | --- | --- | --- | --- |
| EndoAB2 | 0.36064 (Base) to 0.54096 (Apex) | 0.37632 | 0.392 | 0.098 |
| EndoAB1 | 0.36064 (Base) to 0.54096 (Apex) | 0.392 | 0.588 | 0.098 |
| EpiAB | 0.36064 | 0.392 | 0.392 (Base) to 0.588 (Apex) | 0.098 |
| MidAB | 0.36064 | 0.36064 (Base) to 0.392 (Apex) | 0.588 | 0.098 |

##### M-cell Setups

In order to understand the effect of physical characteristics of the M-cells on arrhythmogenesis, the position, shape and size of the M-cells are altered in different setups.

- *Size:* M-cell island is modelled as an ellipse positioned vertically with its center located in the mid layer at position (35,125), and we have considered four different setup and termed as 25M, 50M, 75M and 100M respectively. Different sizes of the M-cell island are considered in which the minor axis is a constant of 10 pixels and the major axis is varied from 25 pixels to 50,75 and 100 pixels respectively. The 100M setup is shown in Fig. 1(i).
- *Position:* The position of the M-cell island is changed by moving the elliptical island of major axis 75 pixels and minor axis 10 pixels to top position (35,75) and bottom position (35,175).
- *Shape:* The shape of the M-cell island is altered by increasing the minor axis to 15 pixels and the major axis is again varied from 25 pixels to 50, 75 and 100 pixels respectively. These setups are termed as Horz-25M, Horz-50M, Horz-75M and Horz-100M.
- *Apex to Base variations:* The apex to base variations which was till now only included in the endo layer in now either introduced in the epi layer or mid layer only. The *G_Ks_* values considered at base and apex are listed in EpiAB & MidAB configuration of Table.3 respectively. The values in between apex and base are linearly interpolated. These four setups are termed as EndoAB2, EndoAB1, EpiAB and MidAB.
- *Poly M-cell island:* In place of a single M-cell island, two M-cell islands were placed in the mid layer. A combination of different sizes and locations are simulated. The different sizes and locations at which the islands are placed are as follows.

– 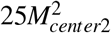: Two M-cell islands with an ellipse of major axis 25 and minor axis 10 located at center (35,25) and (35,225).
– 25*M*^2^: Two M-cell islands with an ellipse of major axis 25 and minor axis 10 located at center (35,75) and (35,175).
– 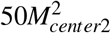: Two M-cell islands with an ellipse of major axis 50 and minor axis 10 located at center (35,50) and (35,200).(shown in Fig. 1(ii))
– 50*M*^2^: Two M-cell islands with an ellipse of major axis 50 and minor axis 10 located at center (35,75) and (35,175). (shown in Fig. 1(iii))
– 75*M*^2^: Two M-cell islands with an ellipse of major axis 75 and minor axis 10 located at center (35,75) and (35,175).
- *M-cell island in endo-mid layers:* Few studies^2^ have reported the presence of M-cell island in the deep subendocardium. Hence, the M-cell island is partially placed in endo-mid layers. M-cell elliptical island with major axis 50 and minor axis 10 is located at center (30,125), and then shifted further into the endo island with center at (25,125). The major axis of the M-cell island is increased by 75 pixels and placed at the same centers. These setups are termed as 50Mendo-1, 50Mendo-2, 75Mendo-1 and 75Mendo-2. The last two setups are illustrated in Fig. 1 (iv-v). In addition, these M-cell island was also moved to the top and bottom position to test its effect on pseudo ECG.

**Figure 1:**
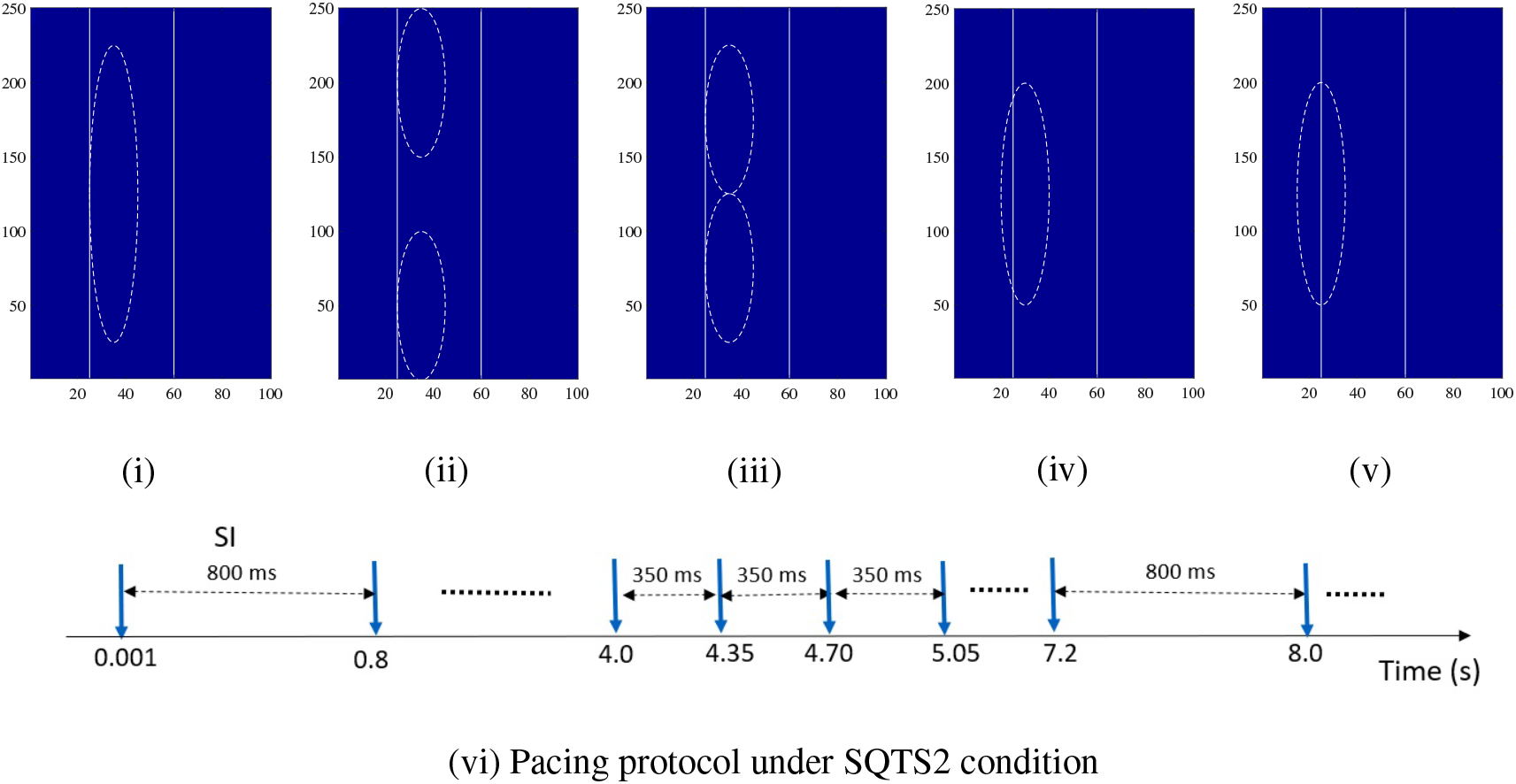
(i) 100M, (ii) 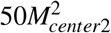, (iii) 50M^2^, (iv) 75Mendo-1 and (v) 75Mendo-2 setups and (vi) Pacing protocol under SQTS2 condition

### 2.4 Model Validation

Inorder to validate which of the setups produced a T-wave that matches with T-wave clinical counterpart, pseudo ECG are synthesized from the depolarization and repolarisation patterns of the model. Further, the performance of the different setups under a disease condition in specific short QT syndrome was analyzed to understand which setups generated an arrhythmia.

#### 2.4.1 Model Validation through Pseudo ECG Synthesis

The ventricular tissue is paced at a cycle length (CL) of 800 ms corresponding to a heart rate of 75 beats/min until all the TP06 variables reach steady state. A cluster of 10×2 endo cells (Cells 1:10,1:2) in the left bottom most corner of the tissue is considered as the normal pacing region. The cells in this region get excited and stimulate the neighbouring cells creating a convex wavefront travelling from the endo to epi layer and from the apex to the base. Pseudo-ECGs are created from the 2D tissue according to the equations 3 and 4 given by Gima & Rudy^25^.

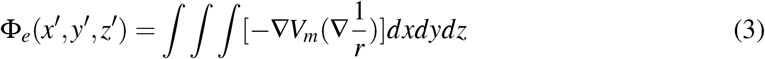

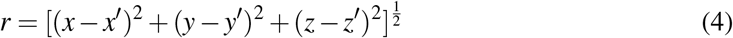

where ∇ *V_m_* is the spatial gradient of voltage and *r* is the distance from a source point (*x,y,z*) to a field point (*x’,y’,z’*). Every point in the gris is considered as a source point while three field points *P*_1_ (95,25,50), *P*_2_ (10,75,50) and *P*_3_ (10,225,50) are chosen on a plane at a distance of 1 cm parallel to the ventricular tissue in the positive z axis to build an einthoven triangle. The voltage difference between *P*_1_ and *P*_2_, *P*_1_ and *P*_3_ and *P*_2_ and *P*_3_ form the three ECG leads. The ECG is normalized and the lead with the highest voltage amplitude (*P*_1_ and *P*_3_) is used for further study. Until now, the composition of the tissue under normal conditions and synthesis of pseudo ECG from this tissue has been described.

#### 2.4.2 Model validation for Specific Condition - Short QT Syndrome

The effect of short QT syndrome type 2 is realized in the model by changing the *I_Ks_* current of all types of cells. The parameters of equations for *I_Ks_* current in the TP06 model are modified based on the experimental data on SQT2 KCNQ1 V307L obtained by Bellocq *et al.*,^28^ Four different cases: wild type(WT), heterozygous (het), homozygous (hom) and homozygous reduced (homred) were considered by Bellocq *et al.*, for KCNQ1 V307L mutation induced changes. Among these, only the heterozygous mutation is considered here as this assumes a 50:50 mixture of WT and mutant KCNQ1 subunits^13^ and has been shown to sustain reentrant arrhythmia. The equation for time constant of *x_s_*-gate of *I_Ks_* current is different for M-cells compared to the other cells since the M-cells have a reduced *I_Ks_* current. These equations are given below.

Heterozygous mutant expression (Het):

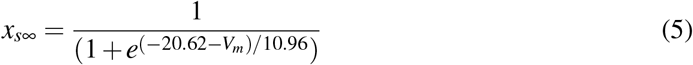

For M-cell

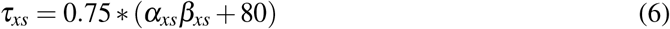

else

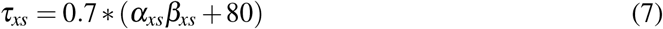

Further, the *I_Ks_* current in epicardial cells is scaled by a factor of 1.5 compared to mid-myocardial cells as reported in the study of Szabo *et al.*, [10], so as to generate a high, symmetrical T-wave as observed in clinical ECG under SQTS conditions. The normal pacing site at the lower left most corner of the tissue is excited at the normal heart rate of 75 beats/min. After all the variables reach steady state, the tissue is now paced at a shorter cycle length (CL) of 350 ms which is representative of premature beats to induce an ventricular arrhythmic pattern. After certain number of such premature beats, the normal pacing CL of 800 ms is resumed. This pacing sequence is shown in Fig.1(vi). The normalized pseudo ECG is then created from the tissue. The tissue activity is recorded for a total duration of 10 s to serve as a digital evidence for cardiac abnormality detection.

## 3 Numerical Results

On simulating the group of cells in the lower leftmost corner (Cells 1:10,1:2) of the tissue, a convex wavefront propagates from the endo to mid and epi layer from the bottom to the top of the tissue. The repolarisation occurs in the epi and endo layer, and M-cells in the mid layer are the last to repolarise. The pseudo ECGs synthesized from the different M-cell setups mentioned in above section 2.3 are described in the following sections.

### 3.1 M-cells as Entire Mid Layer

When the percentage of cells in the different layers is varied for the three different setups such as TM-25, TM-30 and TM-40, the shape of the T-wave is observed to change as seen in the pseudo ECGs in Fig. 2(i). The higher percentage of epi cells in TM-40 is observed to give rise to a low amplitude (0.059 mV and 0.078 mV) notched pattern of T-waves which are usually found to be symptomatic in hypokalaemia condition or long QT syndrome type 2. The peak amplitudes of the T-wave in TM-25 and TM-30 is found to 0.3949 mV and 0.2776 mV respectively. Due to the higher peak value of T-wave obtained in TM-25 setup, the same proportion of cells is used for further analysis. The Tend for the three setups: TM-25, TM-30 and TM-40 occurs at almost the same time instant of 0.41 s, 0.39 s and 0.405 s. The results obtained here for the different percentages obtained for the transmural layers is not in line with that reported earlier by Keller *et al.*,^11^. Their results showed that the TM-40 *A*^1.5^ *B*^1^ setup showed good agreement with the clinically observed results, while in this paper, the same setup, gave rise to a notched T-wave. This may be due to differences in the 2D structure considered as well as a reduction in GJC considered in the mid-epi layer in this study.

**Figure 2:**
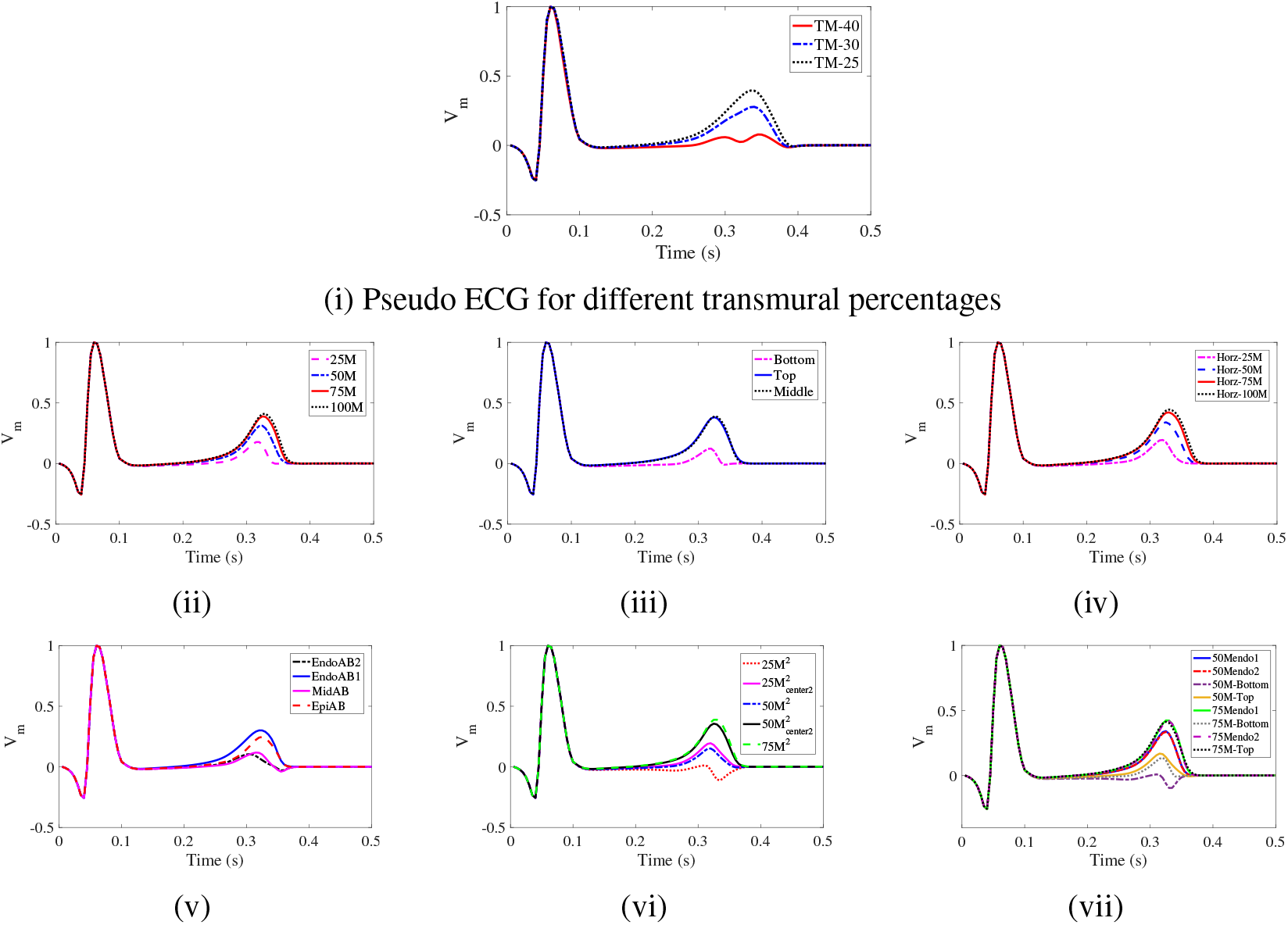
Normalised Pseudo ECGs generated on varying (ii) Size, (iii) Position, (iv) Shape, (v) Apex-Base properties and on introducing, (vi) Two M-cell islands in mid layer and (vii) M-cell in endo-mid layer.

### 3.2 M-cell as Island

The pseudo ECG generated for the different setups of M-cell island explained in section 2.3.2 are shown in fig. 2. It is observed that as the major axis of the M-cell island is varied from 25, 50, 75, and 100, corresponding T-peak value is 0.1771 mV, 0.312 mV, 0.3894 mV, and 0.41 mV respectively. Further, the end of the T-wave is also increased from 0.345 s, 0.375 s, 0.38 s and 0.39 s respectively as seen in Fig. 2(ii). This infers that as the number of M-cells island increases, the time taken for it to completely repolarise increases, and causes the T-end to get lengthened.

When the pseudo ECG patterns generated for the different positions of the M-cell island such as (35,75), (35,125) and (35,175) pixels centered in locations are shown in Fig. 2(iii). Location of M-cell is observed that the T peak is almost similar for the top and middle locations i.e about 0.3826 mV and 0.3894 mV respectively; whereas for the lower position at (35,175), the T-peak value is 0.1233 mV. The T-end values for the three M-cell island positions are 0.385 s, 0.38 s and 0.34 s respectively. Presence of M-cell island in the top or middle position results almost similar T-peak amplitude and duration, while the bottom position have a reduced amplitude and duration. This effect could have occurred due to the depolarisation of electrical impulse propagation pattern is from the bottom to the top of the tissue, this result in exciting the M-cells earlier and return to their resting state also earlier.

When the minor axis of the M-cell island is changed to 15 pixels and the major axis is changed to different values of 25, 50, 75 and 100 pixels, the T-peak value increases gradually from 0.1948 mV, 0.3403 mV, 0.422 mV and 0.4455 mV as seen in Fig. 2(iv). The T-end values are 0.36 s, 0.375 s, 0.38 s and 0.385 s respectively. Similarly, observation of increasing the size of the M-cell island has an effect on increasing the T-peak and T-end values.

In Endo-AB1 setup, a positive T-wave peak of 0.3007 mV is observed. In Endo-AB2 setup, a biphasic T-wave with a positive peak value of 0.1046 mV and a negative peak value of −0.029 mV is represented by the magenta line in Fig. 2(v). This pseudo ECG is characteristic during ischemic condition. If apico-basal variation in included in the epi layer: Epi-AB setup, the pseudo ECG obtained is also concordant although with a lower peak amplitude of 0.2465 mV. However, in the midAB setup with apico-basal variation in the mid layer, a biphasic T-wave with a positive peak value of 0.1179 mV and a negative peak value of −0.03 mV is observed in the ECG similar to that seen in EndoAB2. The T-end is same in all the setups and occurs at 0.375 s. Apico-basal(AB) electrical heterogeneity has been shown to exist in the canine and human ventricular myocardium^27^. In the previous studies, this AB heterogeneity has been included only in the endo layer. In the current study, it is included in the mid and epi layer to understand its effect on the generation of T-waves. Mid AB gives rise to a biphasic T-wave while endo or epi AB gives rise to concordant T-waves. However, M-cell islands in the epicardial positions was not tested or considerd as most of the experimental observations reported that M-cells are presented in the midmyocardium closer to the deep sub-endocardium.

In the 25*M*^2^ setup, a biphasic T-wave pattern with a positive peak value of 0.012 mV and a negative peak value of −0.1116 mV is observed in Fig. 2(vi). The Tend occurs at 0.375 s. When these two M-cell are shifted to different centers in the 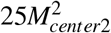, a positive T-peak of 0.1947 mV is seen and Tend is at 0.365 s. The size of the islands are increased to 50 pixels in the 50*M*^2^ setup, here the T-peak value is 0.1517 mV and T-end is 0.36 s. In the 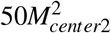 setup, the T-peak value is 0.3563 mV and Tend is 0.38 s. In the 75*M*^2^ setup, the T-peak is 0.3889 mV and T-end is 0.38 s. Thus, when two M-cell islands are placed in the mid layer, it is observed that the T-wave amplitude is higher when the two islands are placed close together than when they are placed further apart.

The placement of M-cell island in the endo layer in both the setups: 75Mendo-1 and 75Mendo-2 has almost similar T-peak value of 0.4185 mV and T-end value of 0.375 s as seen in Fig. 2(vii). When this M-cell island is moved to the top position, the T-peak slightly decreases to 0.4137 mV while the T-end is the same value of 0.375 s. Placing the same island in the bottom position reduces the T-peak to 0.1367 mV and T-end to 0.345 s. Similarly, the reduced size of the M-cell island: 50Mendo-1 and 50Mendo-2 setups also have similar values of 0.3416 mV and 0.3359 mV respectively. The T-end is the same value of 0.37 s for both cases. When these islands are moved to the top position (30,75), the T-peak has a significantly lower value of 0.1692 mV and T-end is 0.365 s. However, placement of M-cell island in the bottom position (30,175) creates a biphasic type of T-wave with positive peak value of 0.0095 mV and negative peak value of −0.095 mV and T-end value of 0.36 s. Current result infers that the T-wave amplitude is increased when the M-cell island is moved from the mid layer slightly into the endo layer, which is inline with Keller *et al.*,^11^ reporting “shifting the M-cells slightly into the endocardial border increases the amplitude of the T-waves”. Further, movement of the island into the endo layer such that the M-cell island is placed in the middle of the endo-mid layer does not alter or change the T-wave amplitude. Later, placement of the M-cell island in the lower position of mid layer creates a T-peak of lower amplitude or a biphasic T-wave. In summary, among the different setups considered, some of them gave rise to concordant T-wave although with different peak amplitudes and duration while few others resulted in a notched or biphasic T-wave. These notched or biphasic waves are generally found in pathologies in which the ventricular repolarisation sequence is altered.

### 3.3 Short QT Syndrome

Based on the different setups of tissue considered for evaluation, only certain setups like TM-25, TM-30, 50M, 75M, 100M, Horz-50M, Horz-75M, Horz-100M, EndoAB1, EpiAB, 75*M*^2^, 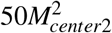, 50Mendo-1, 50Mendo-2, 75Mendo-1, 75Mendo-2, 75M-top generate a positive T-wave peak. Hence, these are considered further for studying which type of tissue is vulnerable to arrhythmia under SQTS2 conditions. Heterozygous mutation, as explained section 2.4.2 was applied in all the cells and the premature beats (PBs) is introduced between the normal pacing pulses to generate an arrhythmia. It was observed, single or two PBs don’t generate an arrhythmia, so 3 PB’s was introduced in-between the normal pacing pulse are and there results are discussed below

#### 3.3.1 M-Cell Layer:TM25 setup

In the TM-25 setup, after six normal pacing pulses of 800 ms, three premature beats of 270 ms are applied in order to generate an arrhythmia. Fig.3(i) shows the pseudo ECG and voltage snapshots on application of this pacing protocol. In the normalised pseudo ECG, it is observed that the first six beats are normal and appear at regular intervals of 800 ms. The peak amplitude of the first beat is 0.4977 mV and the QT interval is 0.305 ms. For instance, Fig.3(ii)(a) shows the voltage snapshot of the pacing site simulated at 0.01 s, a convex depolarisation wavefront originating from the lower leftmost corner of the endo cells is generated that reaches the mid layer at 0.022 s. This wavefront reaches the mid-epi interface after 0.045 s with a delay, due to the lowered gap junction conductances as seen in Fig.3(ii)(b). The depolarisation proceeds towards the end of the epi cells was at 0.07 s and then continues to move upwards. Further, it has been observed that all the cells get depolarised after 0.1 s (not shown here). Repolarisation proceeds from the epi, endo and mid (M) cells from the bottom to the top of the tissue. However, the M-cells located in the top of mid layer are the last to repolarise as seen in Fig. 3(ii)(c) at 0.385 s.

**Figure 3:**
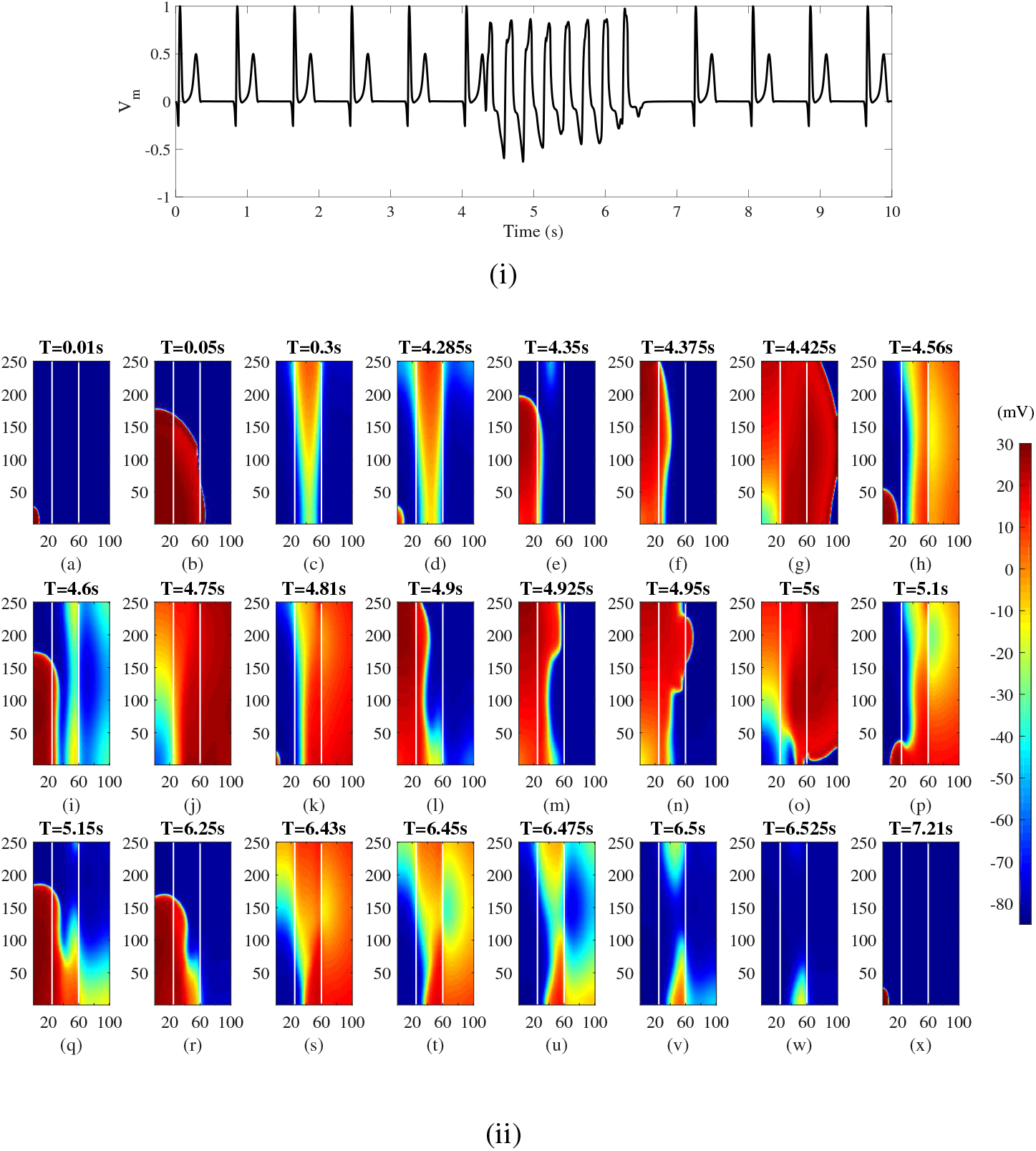
(i)Pseudo ECG and (ii)voltage snapshots in TM-25 setup and heterozygous mutation

Later, three premature beats was applied at 4.27 s, 4.54 s and 4.81 s, after which an arrhythmic type of pattern is observed that is not sustained. After a long pause, the normal pacing pulse is resumed at 7.2 s.

- The First premature beat response is shown in the Fig. 3(ii)(d), where the M-cells in the mid layer of the tissue are still in repolarising state. Therefore, the depolarisation wavefront moves upwards along the endo layer as seen in Fig. 3(ii)(e). Once all the M-cells return to their resting state, the depolarisation wavefront moves from the endo to the mid layer and the epi layer looking similar to a parallel wavefront as seen in Figs. 3(ii)(f-g).
- Fig. 3(ii)(h) shows the state of the tissue after the application of the ‘second premature beat’. It is observed that the cells in the mid and epi layer are in repolarising state. Thus the depoarisation wavefront from this stimulus also travels upwards along the endo layer and then moves into the mid and epi layers as seen in Fig. 3(ii)(i-j) similar to the one seen on the application of the earlier beat.
- Fig. 3(ii)(k) shows the tissue on application of the third premature beat at 4.81 s. The cells in the mid and epi layer are in a depolarised state and therefore the wavefront moves upwards along the endo layer and then reenters into the mid layer and epi layer at the top portion as seen in Figs. 3(ii)(l-n). In Figs. 3(ii)(o-q), it is seen that this depolarisation wavefront travels downward along the epi layer and then reenters into the endo layer. This wavefront continues to circulate within the tissue creating a reentrant arrhythmia as seen in Figs. 3(ii)(r-s) during the time interval till 6.43 s. At 6.45 s, it is observed that the depolarisation wavefront at the bottom mid cells doesn’t reenter into the endo cells as seen in Fig. 3(ii)(t). Therefore, all the cells in the tissue get repolarised as seen in Figs. 3(ii)(u-w).

At 7.2 s, the normal pacing pulse is resumed and this is shown in Fig. 3(ii)(x).

#### 3.3.2 M-cell Layer: TM30 setup

In the TM-30 setup, the heterozygous mutation is incorporated in all cells and the tissue is paced with the pacing protocol as explained in section 2.4.2 section. The generated normalised pseudo ECG is shown in Fig.4. After six normal pacing pulses, three PB’s are applied. Pseudo ECG showed a T-peak is 0.3035 mV and QT interval is 0.30 s, and the three beats give rise to a arrhythmia type pattern; but were not able sustained. Normal pacing sequence was resumed at 5.6 s.

**Figure 4:**
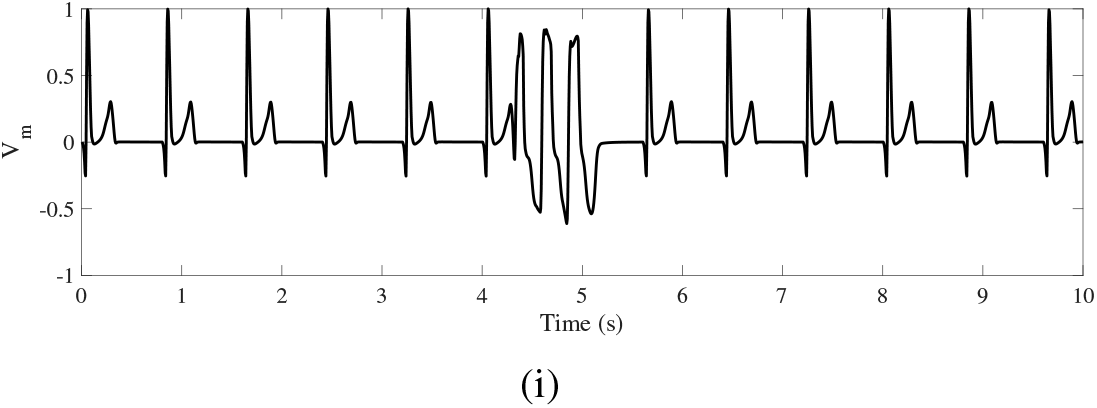
(i)Pseudo ECG and (ii)voltage snapshots in TM-30 setup and heterozygous mutation

#### 3.3.3 M-cell as Island for Short QT Syndrome Condition

In the M-cell Island setup: M75, heterozygous gene mutation of SQTS2 conditions are introduced and the same pacing protocol of premature beats each of 260 ms are applied in between the normal pacing pulses. Fig.5(i) shows the normalised pseudo ECG and voltage snapshots of this M-cell setup on applying this same pacing protocol. In the pseudo ECG, it is observed that the peak value of the first QRS complex is 0.581 mV and QT interval is 0.28 s. After six normal pacing pulses, the three premature beats (PC) are applied at 4.26 s, 4.52 s and 4.78 s. The application of three PB’s doesn’t generate an arrhythmic rhythm and the normal pacing sequence is resumed after a pause at 5.6 s. In Fig.5(ii)(a), the normal pacing site is excited creating a depolarisation wavefront that propagates from the endo cells reaching the mid layer at 0.02 s. At the mid-epi interface in Fig.5(ii)(b), the depolarisation wavefront is delayed due to the lowered gap junction conductance as seen earlier in section 3.3.1. The depolarisation finally reaches the end of the epi layer at 0.07 s and then moves from bottom to the top of the tissue. Repolarisation starts from the epi and the endo layer and the M-cells located as an island in the mid layer are the last to repolarise. Due to the apex-base variation in APD of endo cells, the cells located at the base (top) of the tissue repolarise later than those located at the endo apex (bottom) of the tissue as seen in Fig.5(ii)(c). All cells repolarise at 0.345 s.

- Fig.5(ii)(d) shows the voltage snapshot after the application of the first premature beat, at this instant, the M-cells are still in the repolarising state. By the time the depolarisation wavefront reaches the M-cells, they have returned to their refractory state and are ready to be re-excitable again. Thus, the depolarisation wavefront arising from this stimulus propagates as described earlier although with a small delay through the M-cell island as seen in Fig.5(ii)(e-f). All the cells get depolarised in Fig.5(ii)(g) and repolarisation starts.
- The second premature beat is applied at 4.52 s and the wavefront generated from it is shown in Fig.5(ii)(h). The cells repolarise from bottom to top with the M-cells repolarising last.
- When the third premature beat is applied at 4.78 s, all the cells are in repolarised state and the depolarisation wavefront arising from this PB is seen in Fig.5(ii)(i) at 4.81 s. This wavefront propagates in a regular fashion and all the cells repolarise such that a normal upright T-wave is created as seen in Fig.5(ii)(j-n). Therefore, no arrhythmia is generated and the normal pacing pattern is resumed.

**Figure 5:**
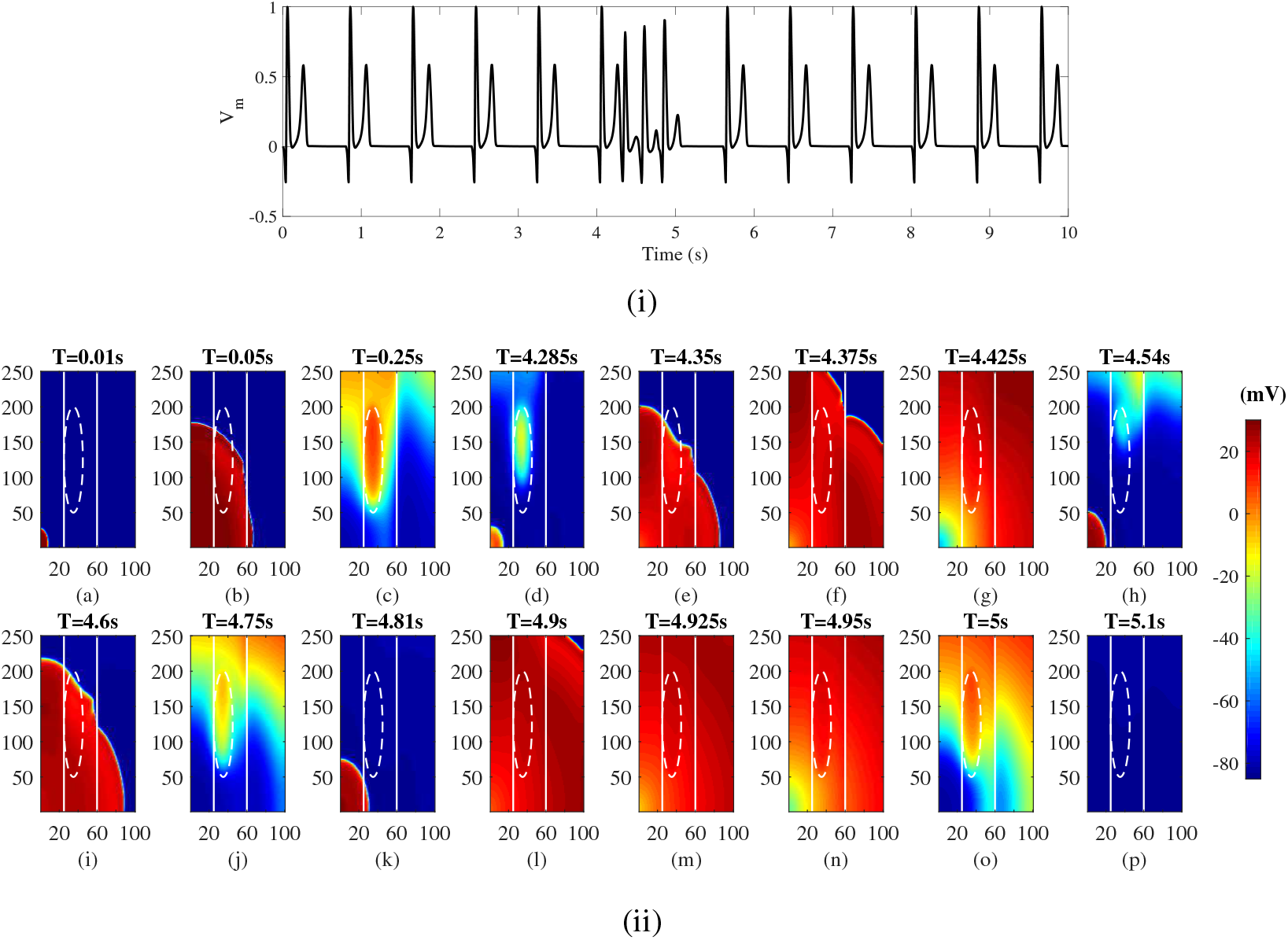
(i) Pseudo ECG and (ii) voltage snapshots in M-cell island setup and heterozygous mutation

The presence of an M-cell island doesn’t generate arrhythmia. This finding contradicts those reported in^16^, where the presence of the M-cell island causes the reentrant arrhythmia to circulate around this island. This could be due to the differences in the electrophysiological properties of the considered M-cell island.

#### 3.3.4 Two M-cell islands

In presence of two M-cell islands having minor axis of 10 pixels, and major axis of 50 pixels located at centers (35,75) and (35,175), the heterozygous gene mutation and same pacing protocol described in section(2.4.2) is applied and the pseudo ECG generated from the tissue is shown in Fig.6(i). It is observed that after the three PB’s, no arrhythmic activity is generated and the normal pacing pulse is resumed at 5.6 s. Further, the T-peak of the first beat is 0.2875 mV and QT interval is 0.28 ms. Fig.6(ii)(a) shows the excitation of the normal pacing site at 0.01 s and the excitation wavefront propagates as usual from endo to epi mid to epi layer as shown in Fig.6(ii)(b). Repolarisation occurs from the epi and endo layers as represented in Fig.6(ii)(c) with the bottom M-cell island repolarising at 0.315 s and finally the top M-cell island repolarising last at 0.345 s. Below are the premature beat observation

- Fig.6(ii)(d) shows that voltage snapshot after application of the first premature beat at 4.26 s. The M-cells located in the top island are still in repolarising state. When the depolarisation wavefront from the first PB reaches the M-cell islands, they have just completed repolarising. Hence, the wavefront passes through the top M-cell island with a small delay as seen in Fig.6(ii)(e-f). All the cells get completely repolarised in Fig. 6(ii)(g) and repolarisation begins.
- Fig.6(ii)(h) shows the voltage snapshot of the tissue after application of the second PB at 4.52 s. The depolarisation wavefront from this proceeds towards mid and epi layer and then repolarisation begins from the epi and endo layers with the M-cell islands repolarising the last as seen in Fig.6(ii)(i-j).
- The third PB is applied at 4.78 s and the depolarisation wavefront from this proceeds in a normal fashion as seen in Fig.6(ii)(k-n). The repolarisation pattern occurs from endo, mid, M and epi cells due to which a biphasic type of T-wave is observed. All the cells get repolarised at 5.135 s.

**Figure 6:**
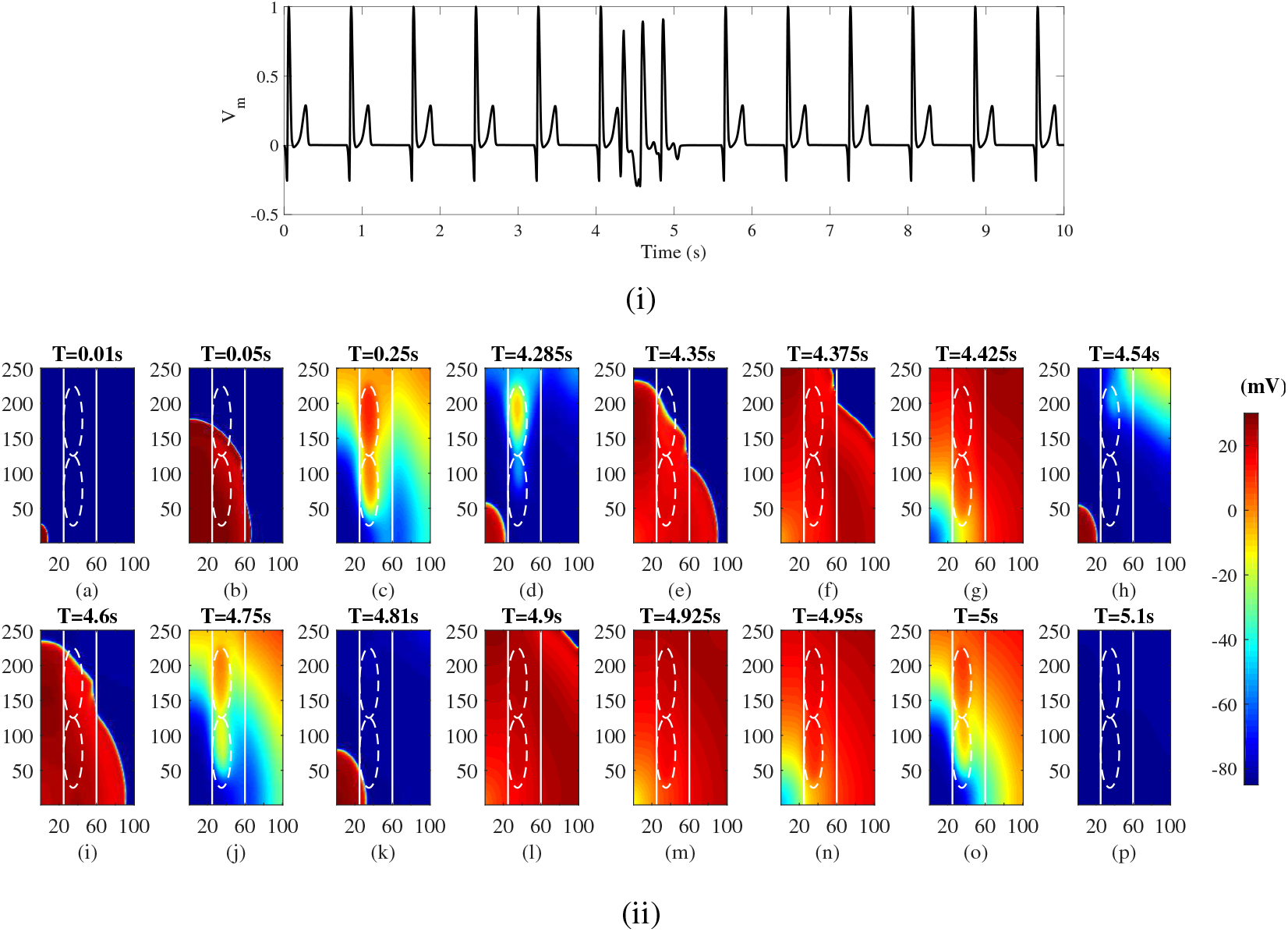
(i)Pseudo ECG and (ii)voltage snapshots in two M-cells setup and heterozygous mutation

Therefore, no arrhythmia is generated.

#### 3.3.5 M-cell in Endo layer

With the M-cell located partially in the endo layer, the same conditions of SQTS2 heterozygous mutation and pacing sequence as in section(2.4.2) are applied. The genearted pseudo ECG in Fig.7 (i) shows that after six normal pacing pulses, the three PBs don’t generate an arrhythmia and the regular pacing sequence is resumed at 5.6 s. The T-peak value is 0.438 mV and QT interval is 0.28 s and the voltage snapshots of the tissue are shown in Fig.7 (ii). The normal pacing site is excited at 0.01 s and the wavefront propagates from the endo layer to mid and epi layer with a delay at the mid-epi interface as seen in Figs.7 (ii)(a-b). Repolarisation occurs from the epi, mid and endo cells with the M-cells repolarising the last as seen in Fig. 7 (ii)(c). This creates a positive T-wave in the first ECG complex. Further, three premature beats are applied and it infers that,

1. The first PB is applied at 4.26 s, and its initially depolarisation wavefront is shown in Fig. 7 (ii)(d). The M-cells located in both the endo and mid layer are still in repolarising state. When the excitation wavefront from the first PB reaches the M-cell island, these cells come into refractory phase and are thus re-excitable again. Thus, the depolarisation wavefront proceeds with a delay through the M-cells as seen in Fig. 7 (ii)(e-f). All the cells depolarise at 4.405 s and is shown in Fig. 7 (ii)(g).
2. The depolarisation wavefront from the second PB is shown in Fig. 7(ii)(h). At this instant the cells in the top are the last to repolarise while the M-cells have already repolarised. This change in the repolarisation pattern creates a biphasic type of T-wave in the first PB. The depolarisation and repolarisation patterns of the second PB are similar to the first PB as seen in Fig. 7 (ii)(i-j) and thus, a similar type of ECG complex is obtained.
3. After the application of the third PB at 4.78 s, the depolarization patterns are shown in Fig. 7 (ii)(k-n). The repolarisation pattern here is different than the earlier two cases with the Mid and epi cells depolarizing last thus forming a slightly positive T-wave. All the cells completely repolarise at 5.135 s.

**Figure 7:**
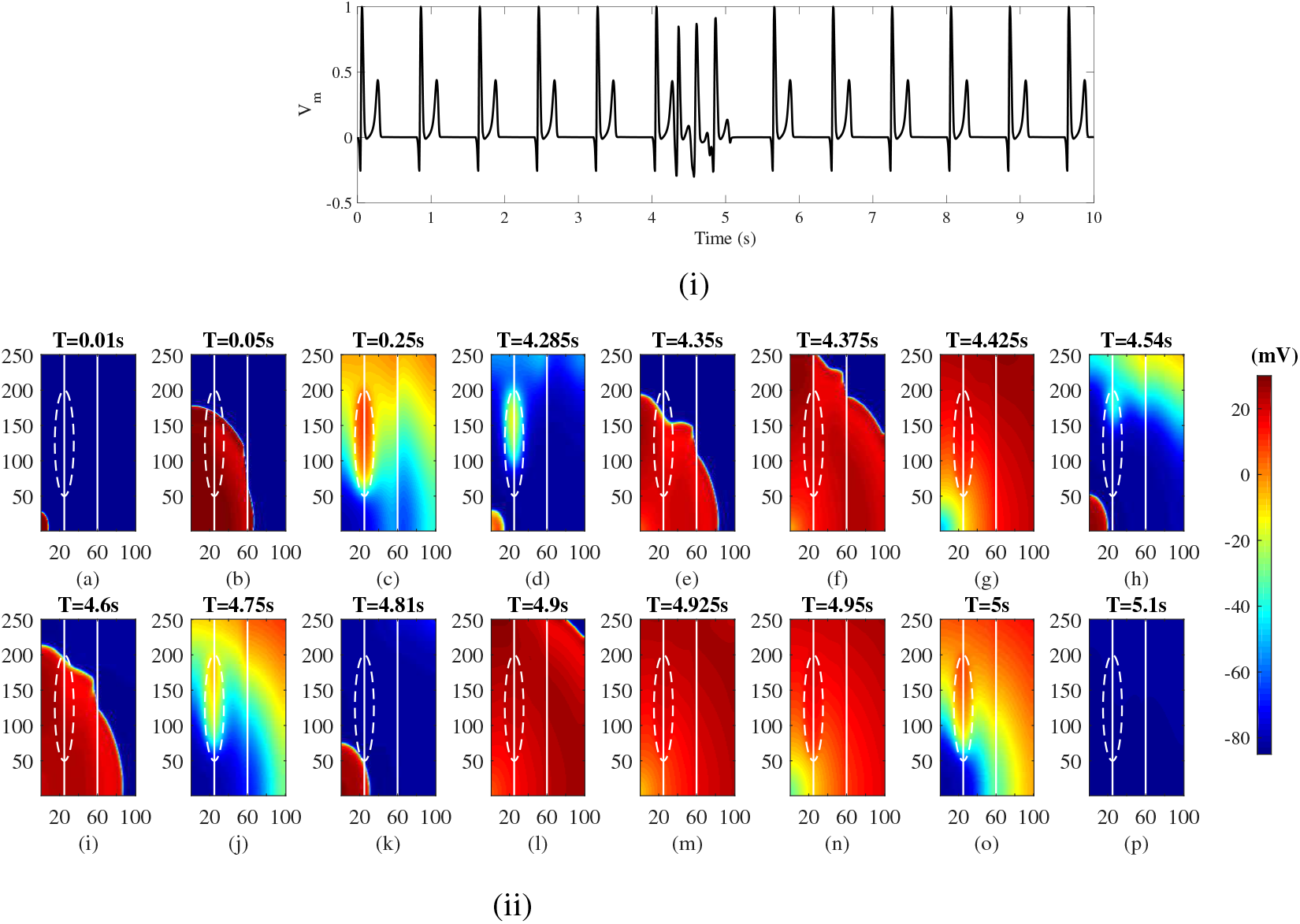
(i)Pseudo ECG and (ii)voltage snapshots in M cell in endo setup and heterozygous mutation

#### 3.3.6 M-cell Island with EpiAB

The apex-base variation in APD which was uptil now only in the endo layer is now introduced only at the epi layer to see its effect on the generation of reentry under SQTS2 conditions. The pseudo ECG in Fig.8(i) shows that on applying 3 PB’s in between the normal pacing pulses, no arrhythmic activity is generated. The T-peak value of the first beat is 0.4574 mV and QT interval is 0.28 ms. The depolarisation pattern generated from stimulating the normal pacing site is shown in Fig. 8(ii)(a-b). Repolarisation is observed to occur from the epi and endo cells with the M-cells repolarising last thus generating positive T-waves for the first QRS complexes. At 4.285 s, the depolarisation wavefront generated on applying the first PB is shown as seen in Fig.8(ii)(c). At this time, the M-cells are still in repolarising state. Thus when the depolarisation wavefront from the first PB reaches the M-cells, there is a delay in the wavefront through the M-cells as shown in Fig.8(ii)(e-f). All the cells depolarise at 4.405 s as seen in Fig.8(ii)(g), the second PB is applied at 4.52 s and its corresponding depolarisation and repolarisation patterns are shown in Fig.8(ii)(h-j); which is similar to the previous PB. At this time, the cells in the top of the mid and epi layer that got excited due to the first PB are the last to repolarise. Thus, a type of biphasic T-wave is observed due to this change in the repolarisation pattern. The third PB is applied at 4.78 s, its depolarisation pattern is shown in Fig.8(ii)(k-n). The repolarisation of the tissue starts in the bottom of the endo layer as seen in Fig.8(ii)(o) followed by the mid, M and epi cells. The epi cells at the top repolarise last. Thus, a low amplitude T-waves is generated in the third PB ECG complex. All the cells are in repolarising state in Fig.8(ii)(p) and finally reach rest state at 5.135 s. Therefore, here also no arrhythmia is generated.

**Figure 8:**
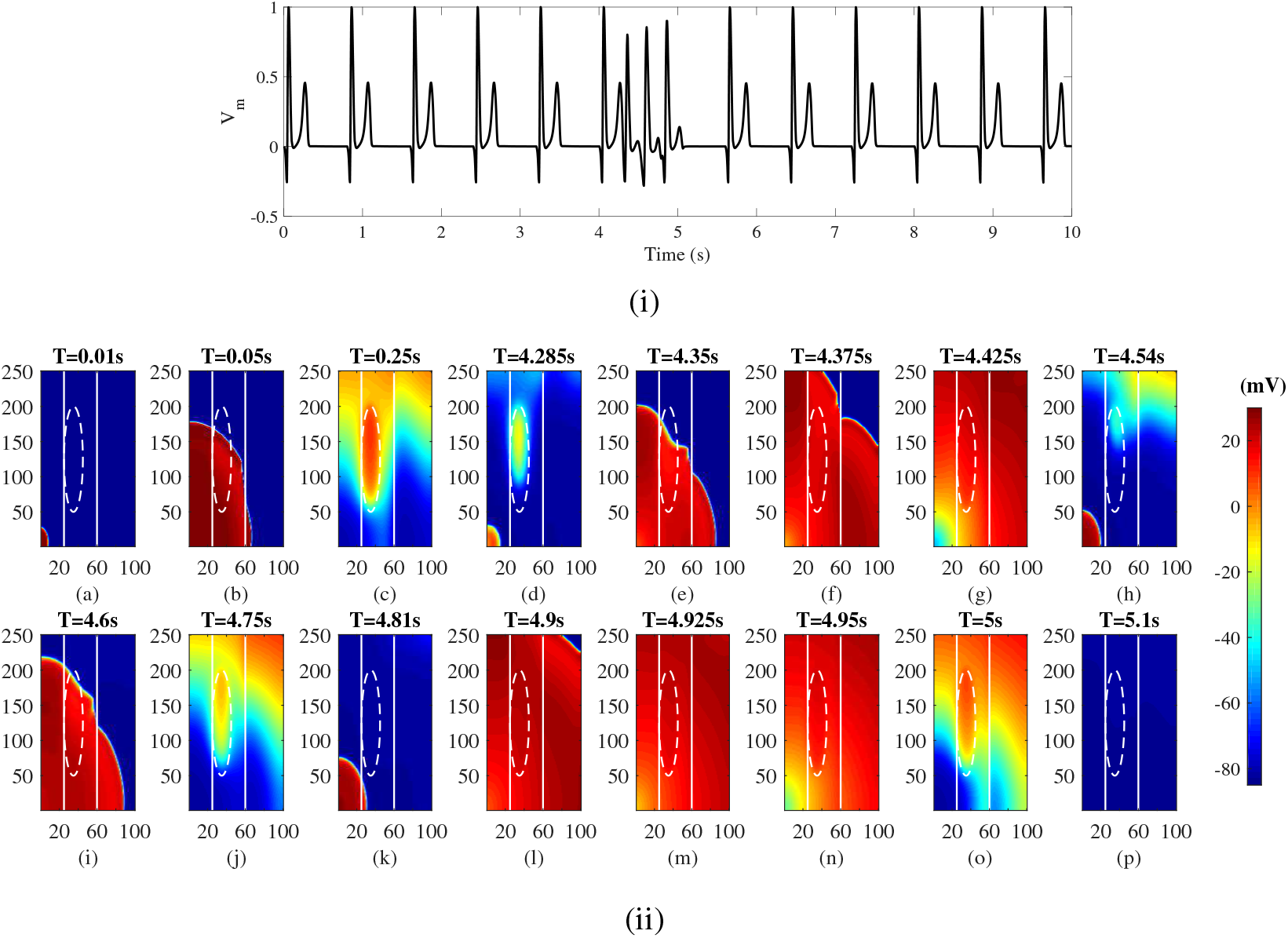
(i) Pseudo ECG and (ii) voltage snapshots in epiAB setup and heterozygous mutation

The sequence of repolarisation in the tissue is dependent on the duration and morphology of AP of the different cells as well as the sequence in which the cells have been activated. Therefore, changing the activation sequence would give rise to different repolarisation patterns and thus different T-wave morphologies. Here, the activation pattern that has been observed in experimental studies occurring from the apex to base from endo to mid to epi layers has been adapted. The limitation of this study is the use of a simplified model of the tissue representing the intrinsic differences in the cells along with the anisotropic variation in conduction. The use of an anatomically accurate model which would take into account the fibre anisotropy and rotation could bring the difference in electrical impulse propagation direction and therefore changes in the repolarisation pattern and subsequently in the morphology of the T-wave.

## 4 Conclusion

In this paper, the authors align with the hypothesis of the existence of M-cells in ventricular tissue and acknowledge that it’s physical and functional characteristics in the generation of T-wave morphology and arrhythmogenesis are unclear. To overcome this gap, this study examined the contribution of M-cells under different setups such as change in size and orientation of M-cells (in the form of a layer or islands). The following different setups, TM-25, TM-30, 50M, 75M, 100M, Horz-50M, Horz-75M, Horz-100M, EndoAB1, EpiAB, 75*M*^2^, 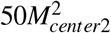, 50Mendo-1, 50Mendo-2, 75Mendo-1, 75Mendo-2, 75M-top showed comparable results with that of its clinical counterpart specifically in generating a concordant T-wave. Thus, this study elucidates the presence of M-cells and it’s important role in characterisation of the T-wave in normal cardiac morphology, when the M-cells are configured as an entire layer or islands in ventricular myocardium. Further, these above mentioned setups were chosen to explain the mechanism of M-cells potential contribution in the generation of arrhythmia in heterozygous SQTS2 condition. Results infer that when the entire mid layer is made up of M-cells, the tissue is vulnerable to arrhythmia under heterozygous SQTS2 gene mutation and when paced with premature beats. On the other-hand, when the M-cells are located in the form of an island, the tissue doesn’t generate any arrhythmia under heterozygous SQTS2 gene mutation.

## Acknowledgements

Funding: This work was supported by Tata Consultancy Services Limited.

